# POPSICLE: A Software Suite to Study Population Structure and Ancestral Determinants of Phenotypes using Whole Genome Sequencing Data

**DOI:** 10.1101/338210

**Authors:** Jahangheer S. Shaik, Asis Khan, Michael E. Grigg

## Abstract

The advent of new sequencing technologies has provided access to genome-wide markers which may be evaluated for their association with phenotypes. Recent studies have leveraged these technologies and sequenced hundreds and sometimes thousands of strains to improve the accuracy of genotype-phenotype predictions. Sequencing of thousands of strains is not practical for many research groups which argues for the formulation of new strategies to improve predictability using lower sample sizes and more cost-effective methods. We introduce here a novel computational algorithm called POPSICLE that leverages the local genetic variations to infer blocks of shared ancestries to construct complex evolutionary relationships. These evolutionary relationships are subsequently visualized using chromosome painting, as admixtures and as clades to acquire general as well as specific ancestral relationships within a population. In addition, POPSICLE evaluates the ancestral blocks for their association with phenotypes thereby bridging two powerful methodologies from population genetics and genome-wide association studies. In comparison to existing tools, POPSICLE offers substantial improvements in terms of accuracy, speed and automation. We evaluated POPSICLE’s ability to find genetic determinants of Artemisinin resistance within *P. falciparum* using 57 randomly selected strains, out of 1,612 that were used in the original study. POPSICLE found *Kelch,* a gene implicated in the original study, to be significant (p-value 0) towards resistance to Artemisinin. We further extended this analysis to find shared ancestries among closely related *P. falciparum, P. reichenowi* and *P. gaboni* species from the *Laverania* subgenus of *Plasmodium.* POPSICLE was able to accurately infer the population structure of the *Laverania* subgenus and detected 4 strains from a chimpanzee in Koulamoutou with significant shared ancestries with *P. falciparum* and *P. gaboni.* We simulated 4 datasets to asses if these shared ancestries indicated a hybrid or mixed infections involving *P. falciparum* and *P. gaboni*. The analysis based on the simulated data and genome-wide heterozygosity profiles of the strains indicate these are most likely mixed infections although the possibility of hybrids cannot be ruled out. POPSICLE is a java-based utility that requires no installation and can be downloaded freely from https://popsicle-admixture.sourceforge.io/

**Author Summary:** The associations between genotypes and phenotypes have traditionally been performed using markers such as single nucleotide polymorphisms. Often, these markers are independently evaluated for their association with phenotypes. A genomic region is deemed significant if multiple markers with significance colocalize. However, multiple markers that are in linkage disequilibrium can sometimes work synergistically and contribute to phenotypic variations. These synergistic associations across markers and across subpopulations have traditionally been captured by population genetic approaches that determine local ancestries. We sought to bridge these two powerful but independent methodologies to improve genotype-phenotype predictions. We developed a new software called POPSICLE that employs an innovative approach to determine local ancestries and evaluates them for their association with phenotypes. Validity of POPSICLE in determining the genes that are responsible for *Plasmodium Falciparum’s* resistance to Artemisinin and in determining the population structure of *Laverania* subgenus of *Plasmodium* are discussed.

## Introduction

Population genetics has traditionally been used to construct admixtures and to reveal shared ancestries genome-wide using tools such as Structure, Fine structure, Admixture, and POPNET (1-14). Some of these tools leverage the common alleles inherited, or co-ancestry matrix to find global ancestries while some others leverage markers such as single nucleotide polymorphisms (SNPs) that are in linkage-disequilibrium to find blocks of shared ancestries within populations. The genome-wide association studies (GWAS) on the other hand were conducted by individually evaluating the markers such as SNPs (15,16). These methods demand hundreds and sometimes thousands of strains to improve the predictability (17). If multiple markers that are deemed significant colocalize, those regions are considered significant in association of genotypes with phenotypes. However, the markers individually may not be significant, but may work synergistically towards contributing to the phenotypic variations. Therefore, there is a need for development of a tool that integrates methodologies from population genetics with those from GWAS. Such a method not only identifies individual markers but also markers in linkage disequilibrium that work synergistically and using lower sample sizes making it attractive to research labs that cannot afford to sequence thousands of strains. We sought to bridge these two powerful but independent methodologies to improve genotype-phenotype predictions. POPSICLE is a novel ancestry determination algorithm that determines the ancestries and leverages them for their association with phenotypes. It offers several improvements in comparison to the existing algorithms. Firstly, conventional algorithms cluster the data by transforming the allele information across the samples into some form of numeric data such as common alleles inherited or allele frequencies within populations, all of which can be found after determining the population size (“K”). Ancestry models for different “K” are evaluated based on how well the data fits the model. This requires running the entire ancestry pipeline for different values of “K”, thus imposing a high computational burden. POPSICLE follows a unique input coding scheme that leverages allele frequencies to find the optimum “K” using cluster evaluation metrics. This allows finding “K” prior to determining ancestries thereby reducing computational time. It also permits incorporating missing data without having to artificially impute the missing alleles or remove useful markers. Secondly, the genome subsequently is divided into blocks of a certain size and the haploblocks are assigned to one of the “K” subpopulations to construct local ancestries. POPSICLE offers flexibility by not forcing the haploblocks into one of the “K” populations. Within a haploblock, not all “K” populations might be relevant as two or more subpopulations might have shared ancestries or more than “K” populations might be needed as some strains might have haploblocks inherited from strains not included in the cohort. Finally, inferring genotype-phenotype associations using local ancestries allows for the assessment of multiple markers that may work synergistically (18,19).

We tested the validity of POPSICLE using 57 (3%) of the 1,612 strains employed in the original study to detect genes implicated in resistance of *Plasmodium falciparum* (*P. falciparum*) to Artemisinin (ART) (17). The *P. falciparum* infections to humans are a relatively recent phenomenon (past 1000 years). The administration of drugs such as Artemisinin(ART) have been used since 1980s to combat the disease (20). Although ART is still one of the first lines of treatment against severe and uncomplicated malaria, various species of *Plasmodium* are increasingly growing resistant to this drug. Several ART derivates have therefore been designed to address the issue of resistance of *P. falciparum* to the original ART. Despite this, the rate at which the ART derivatives have cleared malarial parasites from the blood has gradually declined in Southeast Asia. This impends the control strategies and overall efficacy of ART combination therapies thereby posing the possibility of its spread to the neighboring continents. To make the matters worse, *P. falciparum* parasites have developed resistance not just to ART and its derivatives, but also to almost all previously administered antimalarial drugs such as chloroquine and their derivatives (21).

We also employed POPSICLE to study the population structure of *Laverania* subgenus of *Plasmodium* to which *P. falciparum* belongs. The causal factors for speciation, adaptations, infectivity and resistance to drugs are poorly understood aspects of malaria research. The unicellular protozoan parasite *P. falciparum* is the deadliest species of *Plasmodium* that infects humans and causes over a million deaths per year (22). *P. falciparum* is prevalent in sub-Saharan Africa (23) and is responsible for 200 million clinical cases and 300000 malaria related deaths annually, predominantly in children under the age of 5 (world health organization, 2015). The evolutionary origins of human infecting *P. falciparum* indicate that they have evolved through a cross-species transmission from African apes. Genome composition indicates that *P. reichenowi*, a parasite that mainly infects chimpanzees is highly similar to *P. falciparum*. Another *Plasmodium* parasite *P. gaboni*, also found in chimpanzees is morphologically similar to *P. falciparum* raising the possibility of a diverse parasitic species from which *P. falciparum* could have evolved (24). Due to high sequence similarity, *P. falciparum, P. reichenowi* and *P. gaboni* are placed under the subgenus *Laverania*. Although *P. falciparum* shows high similarity to *P. reichenowi* and *P. gaboni*, it is suspected to have little to no intra-species genetic diversity whereas *P. reichenowi* and *P. gaboni* show a greater diversity (up to 10 times more) (25,26). The ability of POPSICLE to detect genes associated with *P. falciparum’s* resistance to ART and its utility in revealing the population structure of *Laverania* subgenus are discussed next.

## Methods

### POPSICLE employs a unique coding scheme that allows finding optimum “K” independent of ancestry determination

POPSICLE employs a unique coding scheme that leverages allele frequencies to determine ancestries. In a population containing multiple haplotypes, the organism’s alleles and their copies play a key role in determining its survival, adaptation, multiplication, and phenotypic diversity. However, most algorithms that determine admixtures leverage allele composition but not their copy numbers. They code the data as “A” if it matches the reference and B if it matches the alternate allele. This and similar coding schemes capture information related to the alleles, but not their frequencies. To address this issue, POPSICLE (Figure 1A) accommodates allele frequencies by using a three-dimensional matrix of markers (M), samples (S) and nucleotides (4: corresponding to A, T, C and G). For example, 0:0.5:0.5:0 means that at a given marker, two nucleotides T and C are each present at 50% frequency respectively. For most populations, there might only be two primary haplotypes but multiple copies of each of them. But, we have coded the data as MxSx4 to accommodate mutations independently acquired by haploblocks to capture heterogeneity in the population. Reference sequences are usually consensus sequences where heterozygous loci are collapsed to homozygous alleles by replacing the heterozygous allele with the most common allele in a population or by random selection. The proposed coding scheme for the reference sequence is flexible enough to code heterozygous loci, which are sometimes represented using IUPAC nucleotide codes in the consensus sequences. The strains in the coded format are compared against the reference and the markers in concordance with the reference are set to 0 and the rest are scored based on their deviation from the reference (Figure 1B). POPSICLE clusters the strains for various “K” using clustering algorithms such as K-means (Figure 1C). The choice of optimum “K” is determined by employing the cluster evaluation metrics such as Dunn index (27-29). After fixing the value of “K”, POPSICLE runs the ancestry determination algorithm just once to facilitate better use of computational resources and therefore reducing the computational time (Figure 1D).

**Figure 1.**
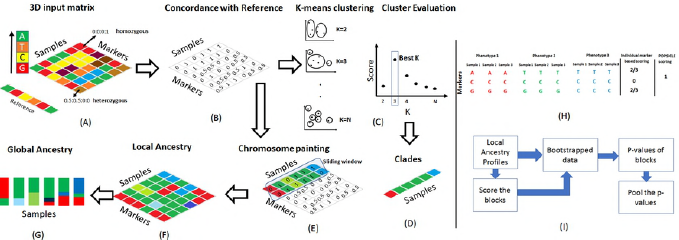
POPSICLE pipeline to infer ancestries and determinants of phenotype. A) The markers are coded using a 4-dimensional vector representing A, T, C and G compositions. Two representative markers are shown in the figure, one homozygous for G is coded (0:0:0:1) and the second heterozygous “AT” is coded 0.5:0.5:0:0. This coding format accommodates copy number variations, aneuploidies and polyploidies. For example, a region with copy number 3 with composition “AAT” can be coded as 0.67:0.33:0:0 allowing incorporation of allele frequency information. B) The markers in concordance with the reference are set to 0 and the rest of the markers are scored based on deviation from reference. This unique coding scheme allows for clustering the data without the need for finding common alleles inherited, or co-ancestry matrix. Data is clustered for various values of “K” and the output clusters are evaluated using metrics such as Dunn Index D) The “K” with the best score is selected and the samples are assigned to different clades. The entire clustering and clade assignment process is independent of ancestry determination thereby improving computational efficiency. E) The input matrix is divided into non-overlapping blocks of user-defined size and the haploblocks are clustered. The strains are assigned to the existing clades or new clades based on the haplogroups. F) The chromosome painting reveals the shared ancestries across the genome. G) This information can be summarized to reveal global ancestries. H) Block-based GWAS based strategy has advantages in comparison to individual marker-based strategy. The markers may not correlate individually with the observed phenotypes and are therefore scored poorly. The block-based strategy on the other hand reveals that the ancestries as represented by distinct colors correlate with the phenotypes and are scored appropriately. I) POPSICLE employs a bootstrapping procedure to verify ancestries against the phenotypes. It creates several bootstrapped datasets and evaluates ancestral blocks for their association with phenotypes in each bootstrapped data. The *p*-values associated with each block are pooled from all the bootstrapped datasets and consensus *p*-values are generated.

### POPSICLE determines global and local ancestries using chromosome-painting like approach

POPSICLE divides the genome into non-overlapping sliding windows of user defined size (e.g. 1kb) and assigns the local profiles to the existing clades or the new clades as needed. This improvement is significant because unlike the existing algorithms, POPSICLE does not force the association of haploblocks to the global clades. Haploblocks can have shared ancestries and therefore, multiple clades can be collapsed into one clade, or new clades might be needed to represent new populations (Figure 1E). These new clades are represented using colors other than the colors associated with the original clades in POPSICLE. The local ancestry determination is repeated by sliding the window across the genome followed by chromosome painting based on the observed ancestries (Figure 1F). The local ancestry patterns reveal genomic segments that are shared, unique segments that are acquired from new populations and the overall genomic divergence. The proportion of different populations present in each strain can be summarized to reveal global ancestries (Figure 1G). Local ancestries can be compared against the observed phenotypes using a bootstrapping approach to find genomic blocks containing one or more markers that are significantly contributing to the phenotypic variations (Figure 1F).

### POPSICLE leverages the ancestries to determine genotype-phenotype associations

POPSICLE employs a block-based strategy that offers significant advantages because individual markers may or may not contribute to the differentiation of the phenotypes but may do so collectively. For example, the three markers in the illustrated block (Figure 1H) do not offer complete association of the genotypes with the phenotypes individually and are therefore scored poorly by traditional methods. The block-based strategy reveals that the genotypes (ACG, TCG and TCC) are consistent in their association with the respective phenotypes and are scored appropriately. The markers that are in linkage-disequilibrium are therefore evaluated and scored based on their intra phenotype-genotype consistency and inter phenotype-genotype differences (Figure 1I). These scores are rank ordered and 60% of the blocks with the bottom scores are putatively considered to follow the null-hypothesis. Since majority of the genes do not contribute to the phenotypic variation, we conservatively used 60% of the blocks as representative of regions that are insignificant. Here, the null-hypothesis is the genotypes do not significantly contribute to phenotypic diversification (30). For each bootstrapped dataset, let “*S*_*b*_” represent the score of the block “*b*” and “S” represent the scores of the blocks that follow the null-hypothesis. The probability value of each block is determined using 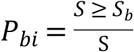. Here, “*i*” is the bootstrap iteration. Assuming “*n*” bootstrap iterations, “n” probability values corresponding to a block are pooled using the Logit method 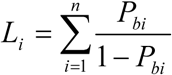(31).

## Results

We sought to evaluate the efficacy of POPSICLE for identifying Artemisinin resistance mutations that confer drug resistance among circulating strains of *P. falciparum* from a large multi-center genome-wide association study that was previously performed to identify genes responsible for resistance to ART (17). The published study used a large sample size of 1,612 samples from 15 regions such as Cambodia, Vietnam, Laos, Thailand, Myanmar and Bangladesh from Southeast Asia, and identified several genes possessing important non-synonymous mutations in coding regions linked to ART resistance. We tested whether POPSICLE could achieve comparable results using a subset of these samples, in this case 57 (out of 1612) samples from 12 (out of 15) locations. Samples were randomly picked manually from the original study with the only selection criteria being that each location contained on average 5 samples (Supplementary File 1). POPSICLE computed the diversity in the selected samples by constructing ancestries and then evaluating how those ancestries contribute to phenotypic diversity. *P. falciparum* are known to exhibit little to no genome diversity (∼1.5 SNPs/kb) thereby posing a challenge to POPSICLE in categorization and determination of ancestries, which directly impacts finding genome-wide determinants to ART resistance.

### Ancestry profile of P. falciparum samples indicates patterns of high genome-wide similarity

The Illumina short reads were aligned to the *P. falciparum* genome 3D7 V.32 (http://plasmodb.org) using BWA and using default parameters yielding an average of 84% alignment rate (Supplementary File 1). We verified if aneuploidies were observed in any of the strains by enumerating the reads that aligned to each chromosome and then translating them into somies. Only minor differences were detected, and these were likely the result of localized copy number variations, which are often detected in gene families (32). No major somy changes suggesting any deviation from haploidy were identified (Figure 2a). Single nucleotide polymorphisms (SNPs) in each of the 57 strains ranged between 50,105 and 1,05,550 SNPs with an average of 73,456 SNPs (supplementary File 1). POPSICLE removed private SNPs and considered each SNP as a separate marker (138029 markers overall). Private SNPs were removed because the alterations specific to individual strains are unlikely to yield phenotypic diversity generally observed across the strains. Allele frequencies were calculated for each of the 57 strains at the SNP markers and to obtain a general overview of the diversity within all samples, a phylogenetic analysis was performed using a hierarchical clustering scheme and complete agglomeration method (33). This indicated that strains from Africa and Bangladesh were the most distinct (Figure 2b) followed by strains from different sub-regions of Thailand and Cambodia. Most of the strains however were assigned to a single branch indicating that the majority of the samples had negligible differences (∼1.5 SNP/Kb). This is consistent with *Plasmodium* literature and supports *P. falciparum* as one of the least divergent species (34). We sought to verify if these strains clustered by geography and found that although some strains such as those from Africa and Bangladesh clustered by geography, the majority of the strains did not. Although the phylogenetic analysis provides a broad overview of the structure of the data, it was not able to identify shared ancestries or where positionally each strain possessed localized differences.

**Figure 2.**
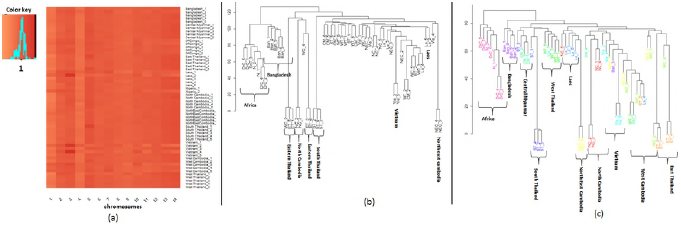
*Plasmodium falciparum* generally show little to no divergence across different strains. The observed somies in *P. Falciparum* indicate that all the chromosomes are haploid although minor variations that may be attributed to factors such as copy number variations are routinely seen, b) Phylogenetic analysis of *P. falciparum generated* using genome-wide markers indicates that except for samples from North Africa and Bangladesh, most samples are similar to each other (∼1.5 SNPs/Kb). c) The genes that varied specifically with respect to geography are determined and phylogenetic tree is drawn using 83 genes deemed significant by POPSICLE. These genes varied significantly by geography, and when used as a feature clustered the strains by geography in the revised phylogenetic tree.

To determine shared ancestries in *P. falciparum*, the POPSICLE pipeline used a block size of 1Kb. A block size of 1kb was chosen because most genes in *P. falciparum* are on average 1kb in size. The choice of other low-resolution blocks such as 5kb or 10kb did not alter the ancestry patterns obtained visually (data not shown). We compared the execution times for POPSICLE with other algorithms such as Admixture and fastStructure (1,2). POPSICLE recorded an execution time of 42 minutes in comparison to 6.82 hours by Admixture and 9.36 hours by fastStructure on a Linux server with standard memory of 100G, virtual memory of 100G and 1 processor (Intel Xeon E5-2670 2.6 Ghz) (Supplementary File 2). The choice of “K” which is obtained by finding population size that offers the least cross-validation error is different for all three algorithms (Supplementary File 2-Figure S1). This is because the ancestry model employed by the individual algorithms dictates the choice of “K”. For the choice of a given “K”, Admixture and fastStructure depicted at least one strain from each clade using a unique color and rest of the strains were assigned colors based on their similarity with different clades. POPSICLE instead works on the philosophy that the samples from a clade are assigned the same color except for the proportion of genome that is different (Supplementary File 2-Figure S2). The ancestry patterns recorded by POPSICLE are more in line with little to no diversification that is commonly observed in *P. falciparum*.

Genome-wide local ancestry patterns revealed that most samples (55%) were assigned to the yellow clade and had only minor variations among them (Figure 3a). These are the samples assigned to the largest branch within the phylogenetic analysis (Figure 3b). However, there were samples that shared ancestries with the yellow clade, but also contained blocks that were clearly different. For example, strains from Africa (DR Congo and Nigeria-blue clade) and South Thailand (green clade) contained many blocks that were clade specific. The samples from Bangladesh, East & West Thailand and North Cambodia also contained many clade-specific blocks. Four of the Ethiopian samples clustered into two clades of two strains each. Clustered into each of these two clades are the two strains from Western Cambodia. The broad differences between these strains, although captured by the phylogenetic analysis, did not show which regions were different genome-wide in the absence of the positional and ancestry information provided by POPICLE. Importantly, the 55% of samples assigned to the yellow clade had only minor variations consistent with a clonal expansion. To verify this further, we examined them at chromosome resolution and found that the yellow clade contained some introgressions with samples from Bangladesh, Africa and South Thailand (Figure 3 b & c)). This argues against clonal expansion theory where highly similar strains may acquire minor independent alterations, but what we saw instead were the shared ancestries among the least divergent and the most divergent strains of *P. falciparum*.

**Figure 3.**
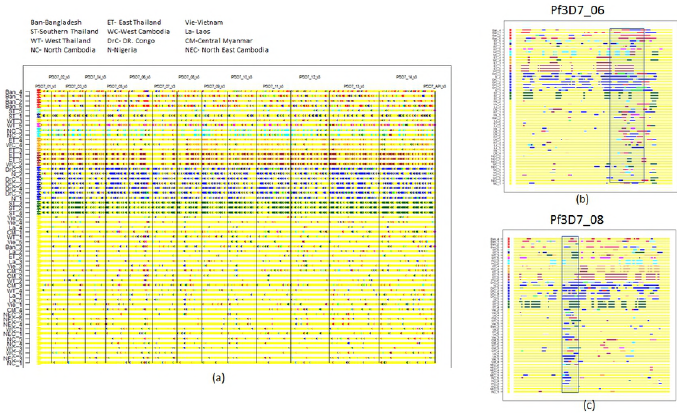
Population structure of *P. falciparum* strains indicate that against a uniform background some differences among the strains are observed. a) The local ancestries using POPSICLE indicate that samples from Africa (DR. Congo, Nigeria) and southern Thailand are the most distinct similar to what is observed in the phylogenetic analysis. The samples from Eastern Thailand, North Cambodia and Bangladesh also show divergence that differentiate them from rest of the samples. Most samples (55%) assigned to the yellow clade were highly similar to each other even though minor variations are observed. b, c) The local ancestries in chromosomes 6 and 8 show blocks of shared ancestries between the yellow clade and the strains from Africa, Bangladesh and South Thailand.

### Some P. falciparum genes evolved rapidly and accumulated geography specific polymorphisms

At chromosome resolution, some strains, irrespective of geography, acquired independent alterations. We therefore queried for broad variations that were geography specific using POPSICLE. Using an adjusted p-value cutoff of 0.05 and restricting the analysis to the SNPs that were intragenic, we found a total of 94 statistically significant blocks (1kb size) that contained 83 genes (supplementary File 3). Although in general, these blocks spanned were found on all chromosomes, chromosomes 1, 7, 8 and 9 were an exception. These represent the chromosomes that did not have any alterations specific to geography. We performed a visual cross-validation of these genes to see if they offered separation of different regions when used as features. These genes are conserved within a region but divergent across different geographical locations and therefore, clustering the samples using these genes should cluster the strains by geography. The phylogenetic analysis performed using the 83 significant genes clustered the strains mostly based on geography, as expected (Figure 2c).

### Genotype-phenotype associations using POPSICLE unravels several key genes/families previously implicated in ART resistance

The original research study employed a large sample set of 1,612 samples extracted from individuals receiving ART treatment for *P. falciparum* infections (17). The parasite loads were recorded every 6 hours and clearance half-life times were documented and used as a clinical phenotype. Using a set of 18,322 SNPs extracted from regions with adequate coverage from 1,063 samples, the original study performed GWAS analysis using a linear regression algorithm implemented in FaST-LMM (35). Based on the missense mutations that had the potential to alter the protein structure, the authors found 9 key genes, 7 with a known function including *kelch, arps10, fd, mdr2, crt* and *pph* and 2 conserved proteins with unknown function that correlated with ART resistance. Although these genes have equal potential towards ART resistance, *Kelch* by far was the gene that was statistically significant containing multiple non-synonymous mutations. We sought to test POPSICLE using a much smaller sample size of 57 to find genes responsible for ART resistance. We employed 1kb blocks and using 100 bootstrapped datasets, found regions that were strongly correlated with the resistance to ART (Figure 4a). We also explored the choice of 5kb and 10kb blocks, which provided us with comparable results. Using p-values adjusted using Benjamini-Hochberg approach and using a cutoff of 0.05, we found a total of 54 1kb blocks that were significantly associated with the phenotype differences (Supplementary File 4). Out of these 54 blocks, 45 were intragenic, which were verified to see if they offered differentiation of the samples based on clearance rates. We categorized the clearance times as slow (>7 hours), medium (between 4 and 7 hours) or fast (less than 7 hours) based on the frequency distribution of the clearance rates and labelled the samples as fast, medium and slow for visualization using principal component analysis (PCA). We performed PCA using the SNP markers that were included in these 45 significant genes. We observed that the samples could be differentiated based on the clearance phenotypes (Figure 4b). We next verified to see if the 9 genes found to be significant in the original research study (17) were present among our list of 45 genes. We found *Kelch* was significantly associated with ART resistance (p-val. 0) but not the other genes from the original study. Since missense mutations have the potential to alter protein structure and therefore function, we found genes on each chromosome that had missense mutations and were deemed significant by POPSICLE. We again found *Kelch* was the most significant gene (p-value 0) with multiple non-synonymous modifications. Other genes such as *rifin, ABC transporter, VAC14* and *Plasmodium exported protein* also contained multiple non-synonymous mutations.

**Figure 4.**
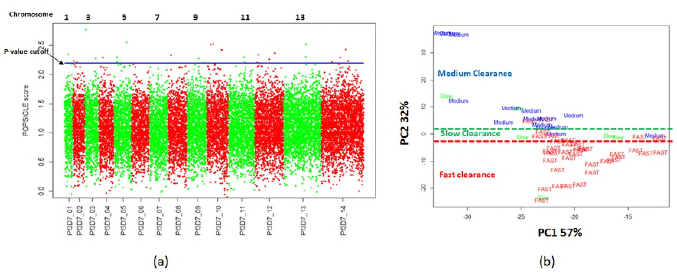
Genotype-phenotype analysis using POPSICLE is performed using 100 bootstrapped datasets and using a block size of 1kb. a) The genome-wide scores for the blocks as determined by POPSICLE are plotted by chromosome. The blue dashed blue line represents the minimum score observed at the chosen adjusted p-value cutoff of 0.05. The blocks with scores above the blue line represent regions significantly correlated with artemisinin resistance. b) Parasite clearance rates are categorized as fast (>7hrs), medium (between 4 and 7 hours), and slow (<7 hours) based on the frequency distribution of the clearance rates. We performed principal component analysis (PCA) using the SNP markers that were included in the 45 genes. We observed that the samples could be differentiated based on the clearance phenotypes using the significant markers from the 45 genes.

### POPSICLE generated ancestral profiles of *Laverania* subgenus indicates coinfections involving *P. falciparum* and *P. gaboni*

Previous studies have indicated genetic similarities between *P. falciparum* and other chimpanzee infecting parasites (34). We therefore studied the population structure of the subgenus *Laverania* using whole genome sequencing data published by multiple research groups (supplementary file 1). The 57 *P. falciparum* DNA extracted from clinical isolates from the ART study were reused for this analysis. The 21 *P. gaboni* samples were subjected to selective genome amplification prior to sequencing and 13 samples (9 *P. reichenowi* and 4 *P. gaboni*) that were sequenced using non-purified DNA were also used. The purity of the sequenced DNA dictates the specificity of the species being sequenced and therefore the overall alignment numbers when aligned to a reference genome of interest. The three species studied are known to show high sequence similarity with high genomic synteny (36). We therefore aligned all the sequences to *P. falciparum* 3D7 genome V.32 (PlasmoDB) using BWA (37). On average, 85% of the reads from *P. falciparum* samples aligned to the genome (Supplementary File 1). The 21 *P. gaboni* samples that were subjected to selective genome amplification showed high alignments (96% on average) whereas the 13 samples sequenced using non-purified DNA showed lower alignments (25% on average).

### Phylogenetic Analysis of *Laverania* subgenus reveals possible coinfections involving *P. falciparum* and *P. gaboni*

We performed the phylogenetic analysis of the *Laverania* subgenus using three distinct sequences of *P. falciparum* apicoplast genome (genbank id: LN999985.1), mitochondrial cyb, cox1 and cox3 (genbank id: M99416.1) and nuclear genome using small subunit ribosomal RNA (SSUrRNA) (genbank id: NC_037282.1). We constructed maximum likelihood trees using Tamura-Nei model and Nearest-neighbor interchange heuristic method implemented in Mega 7 software (38). The phylogenetic analysis using the apicoplast genome revealed 3 major clades corresponding to the three species from *Laverania* (Figure 5A). However, we observed that 4 of the *P. gaboni* samples from the study ERP000135 (36) clustered with the *P. falciparum* and not with the other *P. gaboni* samples. These 4 samples were extracted from a single young male chimpanzee in Koulamoutou, Ogooue-Lolo province in Gabon (36). This suggests that these *P. gaboni* samples are maternally linked to *P. falciparum* rather than with *P. gaboni*. We performed a similar analysis using the genes cytochrome b (Cyb) and cytochrome oxidase subunits 1 and 3 (Cox1 and Cox3), which are part of mitochondrial genome. The phylogenetic analysis revealed 3 major clades, similar to what was observed using apicoplast genome (Figure 5B). The 4 *P. gaboni* samples again clustered with the *P. falciparum* samples confirming similar maternal mitochondrial inheritance from *P. falciparum*. We performed a third phylogenetic analysis using the 18S sequence from the nuclear genome and found 3 clades similar to what was observed using the apicoplast and mitochondrial sequences (Figure 5C). Again, these 4 *P. gaboni* samples clearly show high similarity with the *P. falciparum* genome rather than with the other *P. gaboni* strains. The high similarity of these four *P. gaboni* samples with the *P. falciparum* strains using SSUrRNA and their common maternal inheritance using apicoplast and mitochondrial genomes together suggest that these might be a result of introgressions between *P. gaboni* and *P. falciparum* or are perhaps coinfections. Irrespective, these samples indicate the presence of *P. falciparum* in the chimp. To differentiate between the existence of a mixed infection between *P. gaboni* with *P. falciparum* from that of inter-specific hybrids, we generated ancestral profiles of these samples using genome-wide SNPs detected with respect to *P. falciparum* 3D7 genome V.32 (Plasmodb) using the software samtools (39). The *P. falciparum* samples on average contain 33,367 SNPs in comparison to 233,962 and 435,193 average SNPs in *P. gaboni* and *P. reichenowi,* respectively (Supplementary File 1). We combined the individual SNPs from strains belonging to all 3 species into a set of 1,302,566 markers and used them to generate ancestral profiles using POPSICLE.

**Figure 5.**
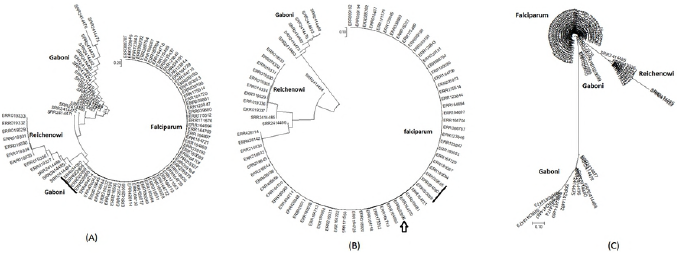
Phylogenetic analysis using apicoplast, mitochondrial (cyb, cox1 and cox3) and small subunit ribosomal RNA sequences. A) Analysis using apicoplast genome suggests three major clades that correspond to the three species being studied. The *P. gaboni* samples extracted from a single chimp from Koulamoutou however clustered with *P. falciparum* instead of being part of the *P. gaboni* clade. B) Analysis using mitochondrial genome markers also supports three major clades, which in general agree with the species assignments. However, similar to the results observed using the apicoplast genome, four *P. gaboni* samples extracted from the same chimp cluster with the *P. falciparum* samples. C) Analysis using the SSUrRNA also clusters those four *P. gaboni* strains with the *P. falciparum* samples instead of the clade consisting of the rest of the *P. gaboni* samples.

### Ancestry profiles of the *Laverania* subgenus reveal patterns indicative of coinfections involving *P. falciparum* and *P. gaboni*

We employed POPSICLE to dissect the population structure of *Laverania* subgenus of *Plasmodium* by unraveling shared ancestries among *P. falciparum, P. gaboni* and *P. reichenowi*. POPSICLE, which leverages K-means clustering assigned these samples to 3 different clades that correlated with the species assignments (Figure 6-Inner concentric circle). The *P. gaboni* samples were assigned to the blue clade, the green clade contained *P. reichenowi* samples and the samples in the red clade corresponded to *P. falciparum*. The majority of the *P. gaboni* strains did not introgress with the *P. falciparum* whereas the *P. reichenowi* strains showed some introgressions with *P. falciparum*. The extent of introgressions between different clades are indicated by the links connecting the clades in the center of the circle. Similar to the results involving apicoplast, mitochondrial and nuclear 18s sequences the four *P. gaboni* strains from Koulamoutou were assigned to the clade containing *P. falciparum* strains. The global ancestry profiles (middle concentric circle) showed extensive introgressions involving *P. falciparum* and *P. gaboni*. The introgressions between *P. falciparum* and *P. gaboni* in the four *P. gaboni* strains were not concentrated in certain blocks nor limited to certain chromosomes, but rather generally found across the genome (Figure 6-outer concentric circle, Figure 7A). Independent analyses of the four samples extracted from the same chimpanzee showed high agreement with each other with similar break points and introgressions. We also combined the reads from these four samples to obtain better coverage and analyzed independently. The ancestries revealed by this sample were similar to what was seen independently in the four *P. gaboni* samples. We further pursued the possibility of this being a mixed infection of more than one species, in the absence of mating. To mimic this, we simulated 4,996,784 paired end reads each of length 100bp using *P. falciparum* and *P. gaboni* V.32 genomes in various proportions (3 datasets) using Wgsim package (39). The read counts were selected such that 40x average coverage was achieved when aligned to the *P. falciparum* genome. We also generated a fourth dataset mimicking a hybrid to pursue the possibility that these four *P. gaboni* strains are hybrids. The mixed infections involved 30:70, 50:50 and 70:30 proportions of *P. falciparum* and *P. gaboni* respectively. We simulated a hybrid by introducing one crossover in chromosomes 1, 5, 6, 8, 11 and 13 and two crossovers in chromosomes 2, 7, 10, 12 and 14 using inhouse scripts and simulating reads using the Wgsim package. The global ancestral profiles (Figure 6-middle concentric circle) suggested that some proportion of genome was inherited from *P. falciparum* and some was inherited from *P. gaboni* but it did not distinguish between mixed infections and a hybrid. The local ancestral profiles (Figure 6-outer concentric circle, Figure 7A) however suggested that the mixed infection involving 30% *P. falciparum* and 70% *P. gaboni* showed a mosaic patterns consistent with what was observed for the four *P. gaboni* strains. Conversely, the simulated hybrid showed clear break points reminiscent of a hybrid (Figure 7B). We next investigated whether heterozygosity was present by quantifying the number of heterozygous SNPs in blocks of 5kb across the genome. Heterozygosity was identified and suggested that the data is more consistent with a mixed infection, rather than an inter-specific hybrid. Specifically, the representative *P. falciparum, P. gaboni* strains, and the simulated hybrid did not show genome-wide heterozygosity (Supplementary Figure 3) whereas the four *P. gaboni* strains from Koulamoutou exhibited genome-wide heterozygosity similar to the simulated mixed infections. Although the proportions of *P. falciparum* and *P. gaboni* in the mixed infection can be debated, based on the ancestral profiles provided by POPSICLE, and the heterozygosity profiles, the data clearly support the four *P. gaboni* strains from Koulamoutou as mixed infections, although the presence of hybrids within these mixed infections cannot be ruled out.

**Figure 6.**
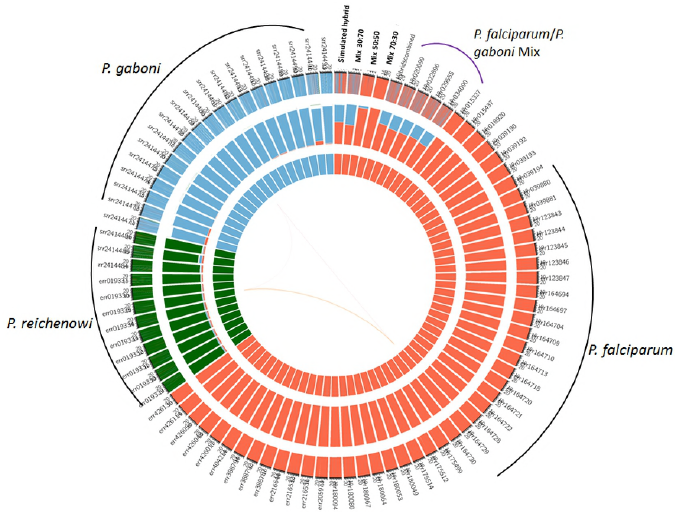
Population structure of *Laverania* subgenus obtained using POPSICLE indicates three major clades that generally agree with the species assignments (inner concentric circle). Exception to the rule are four gaboni samples extracted from Koulamoutou, Ogooue-Lolo province in Gabon that clustered with *P. falciparum*. The global ancestry profiles (middle concentric circle) indicate these samples are either mixed infections or hybrids. To verify if these are hybrids, we simulated three datasets to mimic mixed infections involving 30:70, 50:50 and 70:30 proportions of *P. falciparum* and *P. gaboni* respectively. We also simulated a fourth dataset mimicking a hybrid between *P. falciparum* and *P. gaboni.* The local ancestral profiles (outer concentric circle-Also see Figure 7) indicate patterns similar to mixed infections and not a hybrid, although mixed infections involving hybrids and both the parents cannot be ruled out.

**Figure 7.**
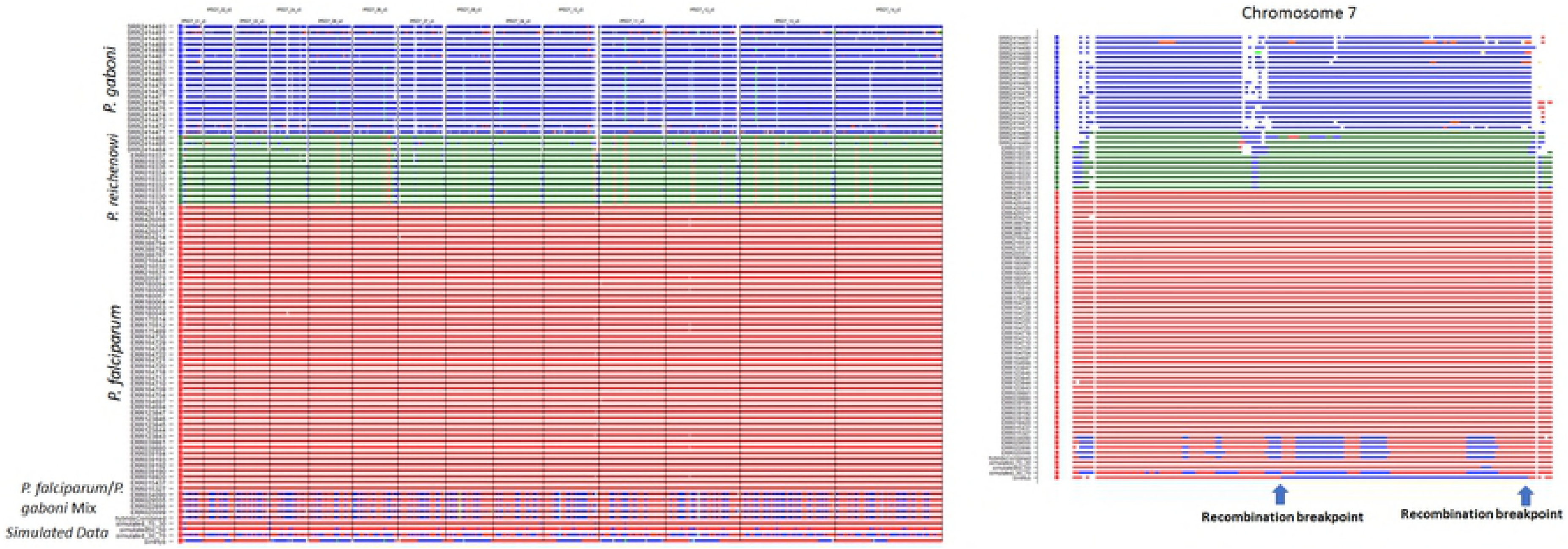
Local ancestral profiles of the *Laverania* subgenus determined using POPSICLE. A) Local ancestral profiles indicate that the clade assignments agree with the species assignments except for four gaboni samples extracted from Koulamoutou, Ogooue-Lolo province in Gabon which clustered with *P. falciparum* instead of *P. gaboni*. Although they clustered with *P. falciparum,* they show intermittent shared ancestries with both the species. The patterns indicate no clear boundaries that are usually expected in the hybrids. To confirm this, we simulated three datasets involving both *P. falciparum* and *P. gaboni* in the ratios of 30:70, 50:50 and 70:30 respectively. We also simulated a fourth dataset mimicking a cross between *P. falciparum* and *P. gaboni.* The patterns shown by the simulated hybrid indicate clear breakpoints similar to what would be expected in a natural hybrid. The patterns shown by 30:70 simulated mixed infections were similar to the observed patterns in the four gaboni samples. B) A high-resolution snapshot of chromosome 7 shows clear break points in the simulated hybrid. The simulated mixed-infections show patterns similar to the patterns from the 4 *P. gaboni* strains from Koulamoutou, Ogooue-Lolo province in Gabon. They show no clear break points and based on the patterns, it is likely that these are similar to a mixed infection (simulated 70:30). We therefore suggest these four *P. gaboni* samples to be mixed infections involving 30-40% *P. falciparum* and 60-70% *P. gaboni,* although mixed infections involving hybrids and both the parents cannot be ruled out.

## Discussion

We introduced a software suite called POPSICLE which unravels the evolutionary relationships within populations and leverages those ancestral relationships to find genomic segments strongly associated with phenotypes. POPSICLE is computationally attractive because it uses a framework that automatically determines the population size prior to ancestry determination. It also allows for the incorporation of missing data without having to impute the missing values, which can lead to errors (4,9). Further, it reveals genotype-phenotype associations using a bootstrapping approach and employs a block-based strategy combining markers that are in linkage disequilibrium. Therefore, it bridges two independent and popular methodologies of population genetics and GWAS. We employed different tests to assess the utility of POPSICLE. We first tested POPSICLE’s ability to find genes associated with ART resistance by employing only 3% of strains used in the original study. The ancestries determined using POPSICLE indicated that *P. falciparum* showed some diversity, although most of the strains were homogenous with little to no diversity. This was not surprising because multiple studies have indicated that *P. falciparum* shows little to no diversity compared to other *Plasmodium* spp., such as *P. vivax,* which possess more than 10-fold diversity (26,34). We observed that 17% of the strains from Africa and South Thailand showed extreme diversity, whereas 28% of strains from Eastern Thailand, North Cambodia and Bangladesh showed less diversity. Majority of the strains (55%) however contained 90% or more shared ancestral blocks irrespective of geographical origin. Although ancestral relations did not correlate with geography for the majority of strains, we uncovered 83 genes that appeared to evolve independently and were significantly correlated with the geography and may reflect specialization for transmission among indigenous vector populations within each geography (supplementary File3). Indeed, many factors such as geography, drug administration, vector species, and complex host-vector interactions likely contribute to the evolution of the parasite. In this multi-factorial process, we honed in on genotypes that correlated strongly with resistance to ART, a drug commonly administered to combat malaria in South-east Asia. Using only 57 randomly selected samples from a cohort of 1,612 samples used in the original study, we were able to find 45 genes associated with drug resistance. *Kelch*, which was most significantly associated with ART resistance in the original study contained multiple non-synonymous mutations and was found to be highly significant in our analysis. In addition to *Kelch*, we also found multiple non-synonymous mutations in the genes for *rifins*, an *ABC transporter, VAC14* and a *Plasmodium exported protein*. Of these, the *rifins* were previously found to be significant to ART resistance in other studies (40-43). Surface antigens such as *rifin, stevor* and *Pfmc-2Tm* have been strongly associated with antigenic variation and the evasion of host immunity (42,43). These surface antigens are proposed to mediate the retention of infected erythrocytes within the microvasculature, blocking blood flow and promoting pathogenesis.

We further employed POPSICLE to unravel cross-species genome-wide similarities among *P. falciparum, P. reichenowi* and *P. gaboni* from *Laverania* subgenus of *Plasmodium. Plasmodium* infections to humans are a relatively recent phenomena and because of the high similarity among the three species of the *Laverania* subgenus, it has long been suspected that *P. falciparum* might have evolved either from *P. gaboni* or *P. reichenowi*, which are known to infect chimpanzees (34). POPSICLE automatically determined the population size to be 3 and the strain assignments agreed with the species designations. Based on the chromosome painting performed by POPSICLE, we discovered four *P. gaboni* samples extracted from Koulamoutou, Ogooue-Lolo province in Gabon that independently showed profiles that appeared to be an inter-specific admixture between *P. falciparum* and *P. gaboni*. Phylogenetic analysis using apicoplast and mitrochondrial genomes of these *P. gaboni* samples supported a maternal origin of *P. falciparum* rather than the expected *P. gaboni* origin. The phylogenetic analysis using nuclear SSUrRNA also indicated the same. We therefore performed independent tests to confirm the nature of these samples. We generated 3 simulated datasets that mimic mixed infections involving 50%-50%, 70%-30% and 30%-70% proportions of *P. falciparum* and *P. gaboni* respectively. In addition, we also simulated a hybrid between *P. falciparum* and *P. gaboni* to verify if the observed patterns in those 4 samples were consistent with the simulated hybrid. The analysis of the mixed-infections is currently only possible with POPSICLE, and at the time of publication, we are unaware of any other tool that leverages allele frequencies to incorporate proportion of alleles present at different markers. Other algorithms simply consider all different proportions of allele frequencies as heterozygous and are thus unable to differentiate between various proportions of mixed infections. The patterns shown by the simulated datasets suggest that the infections in the four *P. gaboni* strains from Koulamoutou were synonymous with a mixed infection involving 70% *P. gaboni* and 30% *P. falciparum.* The patterns in the hybrid simulation, on the other hand, were not mosaic but contained clear break points reminiscent of what would be expected in a hybrid. We further validated our findings by quantifying the heterozygosity in the samples to establish that the four *P. gaboni* strains from Koulamoutou, as well as the simulated mixed infections,showed genome-wide heterozygosity as expected for a mixed infection whereas the representative *P. falciparum, P. gaboni* and simulated hybrid did not. These results argue strongly in favor of a mixed infection rather than the hybrid nature for these four *P. gaboni* strains. This is interesting because these mixed infections involved species that infect humans and chimpanzees. Recently, a research study that investigated the role of insect vectors involved in the inter-organism transfer of parasites found some mosquitoes that were infected with *Plasmodium* parasites from gorillas and chimpanzees (44). They also found that the vectors infected with ape parasites also feed on humans who live nearby and therefore serve as bridge vectors. Since the mosquitoes that usually feed on humans can also bite apes, and vice versa, there is no barrier to transfer of parasites between different hosts. These conclusions argue strongly in favor of the possibility for mixed infections within vectors that feed on different hosts and accumulate multiple *Plasmodium* spp. But since many of the analyses that confirm the species of parasites are based on apicoplast, mitochondrial or SSUrRNA genome, the possibility for the existence of hybrids has not been thoroughly investigated. Although our analysis suggests that the four *P. gaboni* strains are most likely mixed infections, we are unable to rule out the possibility of them being mixtures of hybrids. These results together strongly advocate in favor of POPSICLE’s ability to reveal novel evolutionary relationships and phenotype-determinants using high-resolution next-generation sequencing data. POPSICLE is a platform independent java-based utility that requires no installation and is freely available for download at https://popsicle-admixture.sourceforge.io/

## Figure legends

**Supplementary Figure 1.**
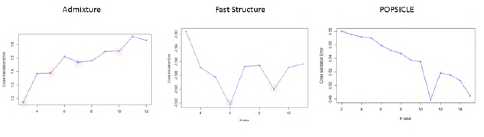
The Choice of “K” is determined by the value of “K” that provides the least cross-validation error. As shown, Admixture’s choice of “K” is 3, Fast Structure’s choice of “K” is 6 and the choice of POPSICLE is 11, which is close to the number of haplogroups present in the data.

**Supplementary Figure 2.**
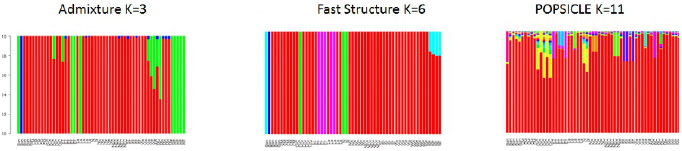
The global admixture patterns as indicated by Admixture suggest three distinct lineages to be present with shared ancestries. Fast structure on the other hand predicts 6 different lineages. POPSICLE suggests admixture that is more in line with what is known about P. falciparum. It clearly shows that although there are variations that can be attributed to some strains, the strains in general are highly similar to each other.

**Supplementary Figure 3.**
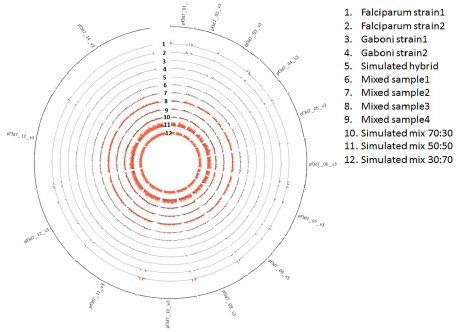
Heterozygosity in various strains as a measure to find mixed infections. Representative *P. falciparum* and *P. gaboni* samples and the simulated hybrid do not exhibit heterozygosity. The four strains (6-9) from Koulamoutou, Ogooue-Lolo province in Gabon on the other hand show extreme heterozygosity. Simulated mixed infections (10-12) also exhibit extreme heterozygosity similar to what is seen in the four samples from Gabon reassuring that these are mixed infections involving *P. falciparum* and *P. gaboni* although the presence of hybrids in this mixture cannot be ruled out.

## Acknowledgments

The authors would like to thank Tracking Resistance to Artemisinin Collaboration (TRAC) and Mahidol Oxford Tropical Medicine Research Unit (MORU) for providing PCt1/2 estimates for the *P. falciparum* study. This study used the Office of Cyber Infrastructure and Computational Biology (OCICB) High Performance Computing (HPC) cluster at the National Institute of Allergy and Infectious Diseases (NIAID). The authors are supported by the Intramural Research Program of the NIAID at the National Institutes of Health, Bethesda, MD.

